# A single latent channel is sufficient for biomedical image segmentation

**DOI:** 10.1101/2021.12.10.472122

**Authors:** Andreas M. Kist, Anne Schützenberger, Stephan Dürr, Marion Semmler

## Abstract

Glottis segmentation is a crucial step to quantify endoscopic footage in laryngeal high-speed videoendoscopy. Recent advances in using deep neural networks for glottis segmentation allow a fully automatic workflow. However, exact knowledge of integral parts of these segmentation deep neural networks remains unknown. Here, we show using systematic ablations that a single latent channel as bottleneck layer is sufficient for glottal area segmentation. We further show that the latent space is an abstraction of the glottal area segmentation relying on three spatially defined pixel subtypes. We provide evidence that the latent space is highly correlated with the glottal area waveform, can be encoded with four bits, and decoded using lean decoders while maintaining a high reconstruction accuracy. Our findings suggest that glottis segmentation is a task that can be highly optimized to gain very efficient and clinical applicable deep neural networks. In future, we believe that online deep learning-assisted monitoring is a game changer in laryngeal examinations.

## Introduction

A functional voice is a crucial factor for a successful social embedding. Voice physiology is commonly qualitatively assessed using stroboscopy [1]. In contrast, laryngeal high-speed videoendoscopy (HSV) is an emerging technique, that also allows quantification of voice physiology [2]. With HSV, the vocal fold motion is typically recorded at several thousand frames per seconds, therefore visualizing each glottal cycle accurately [3]. The glottis, the opening between the vocal folds, is a good proxy for the cyclic behavior and is of major interest for quantitative data analysis.

The glottis can be segmented using several image analysis techniques [4], among others active contours [5], Gabor filters [6] and thresholding combined with level set methods [7]. Recently, deep neural networks for semantic segmentation have been utilized for glottis segmentation [8, 9]. Additionally, optimized deep neural networks for clinical applicability have been proposed [10]. However, these deep neural networks have commonly a black box character, lowering their acceptance in a clinical environment [11]. This effect can be reduced when providing insights to the algorithm. Despite the fact that we know that encoder-decoder architectures and variations thereof are well suited for this task, we are lacking a fundamental understanding of what are the necessities of these deep neural networks.

Autoencoders or, in general, encoder-decoder architectures have contracting and expansion paths [12, 13], where the bottleneck layer is referred to as code layer or latent space. The latent space is thought to contain a high-level embedding of the raw input image. The inspection of this latent space is highly interesting for generative adversarial networks (GANs), as the latent space can be used in GANs to specifically direct the generative image to enable face editing [14, 15], image embedding interpolation [16] and novelty detection [17]. For semantic segmentation, the latent space has also been shown beneficial in multi-task architectures [18, 19]. However, little is known what the latent space represents in biomedical image analysis.

In this work, we are characterizing the latent space, a higher order representation of the endoscopic image, embedded in a semantic segmentation architecture. We systematically investigate its properties and how alterations to the latent space result in differences in the glottis segmentation prediction. With this, we will leverage the potential of latent space information in a clinical context.

## Materials and methods

### Data and preprocessing

To train and evaluate deep neural networks, we used the Benchmark for Automatic Glottis Segmentation (BAGLS, [9]). We used the full training and test dataset containing 55,750 and 3,500 images, respectively. For training, we resized all images to 512 × 256 px, which is the native resolution of most images in the dataset. All endoscopic images were converted to grayscale. The input image intensities were normalized to −1 and 1, the segmentation masks were normalized to 0 and 1, where 0 is background and 1 the glottal area. We applied data augmentation to the training data randomly using Gaussian blur, rotation (−30 to 30°), horizontal flip and gamma adjustments. We also use short video snippets of 30 frames available in the BAGLS dataset for time variant data analysis. Videos are processed as single frames on a single frame basis.

#### Glottal area waveform (GAW)

The glottal area waveform (GAW) is a one-dimensional representation of the vocal fold oscillation behavior. Each time point of the GAW is computed as the sum of foreground pixels in the glottal area segmentation mask at the given time point [20].

### Deep Neural Networks

#### Architecture

The baseline glottis segmentation network is based on the U-Net architecture [21] modified as described in [10]. Briefly, we rely on an encoder-decoder architecture to change the image domain from endoscopic image to glottal area segmentation (see Figure 1). Initially, we use skip connections between encoder and decoder to pass mid-level information by concatenation. We setup deep neural networks in TensorFlow 2.6 using the Keras high-level package. We trained for 25 epochs at a constant learning rate of 10^−4^ using the Adam optimizer [22]. Each convolutional layer used a kernel size of 3×3 and *f*_*L*_ convolutional filters that follow equation 1:

**Fig 1.**
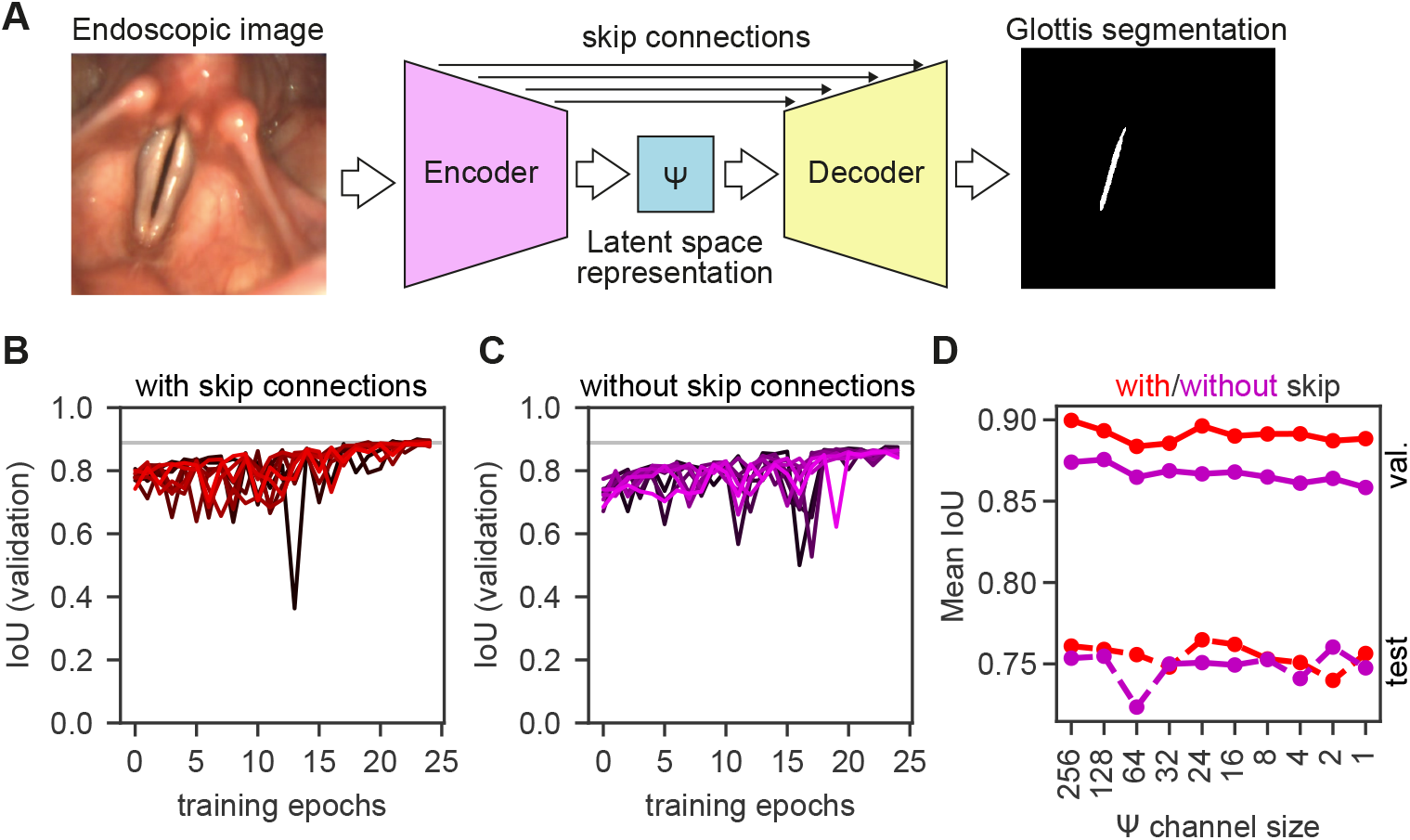
A single latent space channel is sufficient for glottis segmentation. A: Glottis segmentation of endoscopic images using deep neural networks (DNNs) with latent space Ψ. B: Convergence of segmentation DNNs across different latent space channels with enabled skip connections. Gradient from black to red indicate less channels. Gray line indicates maximum IoU score. C: Convergence of segmentation DNNs across different latent space channels with disabled skip connections. Gradient from black to magenta indicate less channels. Gray line indicates maximum IoU score from panel B. D: Performance of best performing segmentation DNNs on validation set (solid lines) and evaluated on test set (dashed lines) with enabled (red) and disabled (magenta) skip connections across latent space (Ψ) channels measured by mean intersection over union (IoU).

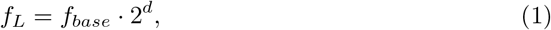

where *f*_*L*_ is the number of convolutional filters which equals to the number of channels in a given layer. At a given network depth *d* (in our baseline model *d* ∈ {0, 1, 2, 3, 4}) and a given initial base filter size *f*_*base*_ (*f*_*base*_ = 16 in our baseline model) we gain a total of *f*_*L*_ = 256 latent space channels at maximum network depth *d* = 4. After each convolution layer, we applied batch normalization. After batch normalization, we used the ReLU function as non-linearity (equation 2). However, in the latent space Ψ (Figure 1A) we used the ReLU6 function (equation 3) that is clipped between 0 and 6:

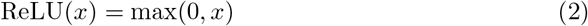

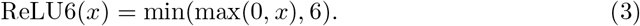

During training, we were minimizing the Dice loss [23] as defined in equation 4 by comparing the predicted glottis segmentation mask *ŷ* to the ground-truth segmentation mask *y*.

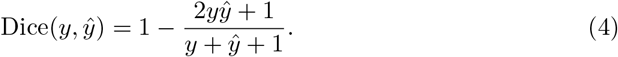

#### Latent space Ψ

The latent space Ψ is a high-level representation of the initial endoscopy image at the end of the encoder and serves as input to the decoder (Figure 1). It can be interpreted as an image with *f*_*L*_ “color” channels. For latent space investigations, we changed *f*_*L*_ from its initial value (here: 256), as defined by equation 1, to a fixed value ranging from 1 to *f*_*L*_. When *f*_*L*_ = 1, we refer to the latent space as latent space image Ψ_1_.

#### Decoder experiments

The initial decoder is constructed as described in the section *Architecture*. For decoder experiments, we used different strategies to construct the decoder (Figure 5A). The latent space image Ψ_1_ is used as sole input to the decoder. Next, we used a combination of Upsampling2D operations and either one or two convolutional layers with *f*_*D*_ channels, where *f*_*D*_ ∈ {1, 2, 4, 8}. For each Upsampling2D-Convolution cycle *UC* ∈ {1, 2, 4}, the Upsampling2D operation uses a scaling factor *s* ∈ {16, 8, 2}, respectively, to ensure a full upsampling to the original image resolution (512× 256 px). For training, we converted each training image to its latent space representation by using the final model used in latent space data analysis (result of Figure 1D, with no skip connections and *f*_*L*_ = 1). We converted each latent space image in uint8 as we have shown that eight bit are sufficient for high-level encoding (Figure 4A,B).

### Bit encoding

For evaluation of the bit encoding, we created a histogram of a given latent space image and divided it into 2^*bits*^ bins. We then set each pixel in a given bin range to the average value in a given bin (Figure 4C). The resulting new latent space image is provided to the decoder and the reconstructed image is compared to the ground-truth segmentation mask. We used the mean squared error (MSE) and the intersection over union (IoU) score (see *Evaluation*) as evaluation metrics.

### Evaluation

We evaluated the segmentation quality using the IoU (intersection over union) score [24] as defined in equation 5.

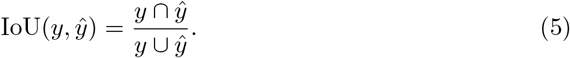

We further computed the correlation between the latent space image Ψ_1_ (in equation refered as *x*) and the GAW (*y*) as follows:

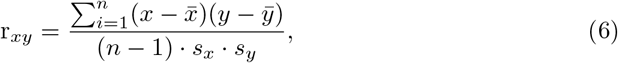

where *n* is the number of time points/samples,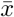 and 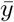 the average of *x* and *y*, respectively, and *s*_*x*_ and *s*_*y*_ the sample standard deviation of *x* and *y*, respectively.

### Code and data availability

We provide all relevant code at https://github.com/ankilab/latent. In this study, we relied on the open BAGLS dataset. We provide all latent space images for decoder training and the used model for latent space image analysis at https://zenodo.org/record/5772799.

## Results

### A single latent space channel is sufficient for glottis segmentation

To understand which components are crucial in a segmentation deep neural network, we performed an ablation study on a modified U-Net architecture (see Methods). We trained a full size, complete U-Net to perform glottis segmentation (Figure 1A), similar to previous works [9, 10]. The latent space Ψ, the ultimate bottleneck that connect encoder and decoder in the full U-Net, has initially 1024 channels (Figure 1A), when 64 filters are used in the first layer (*f*_*base*_=64). In this work, we use a reference implementation with 16 filters in the first layer (*f*_*base*_=16) and thus, 256 channels in the latent space, as this has been shown previously to provide comparable performance compared to *f*_*L*_ = 1024 [10]. We systematically reduced the amount of channels in the latent space to determine the minimum viable latent space. We found that even a single latent space channel is sufficient to encode the glottal area segmentation (Figure 1B). However, we hypothesized that the skip connections in the U-Net allow to rescue the strong limitation in the bottleneck. Hence, we removed the skip connections and found that the segmentation accuracy was reduced across configurations (Figure 1C). However, the network architecture is still able to provide accurate glottis segmentations (Figure 1D), and has a performance on the test set similar to higher latent space encodings and enabled skip connections. In summary, we show that a single latent space channel is sufficient for glottis segmentation. We will refer to this single latent space channel as latent space image Ψ_1_.

### The latent space encodes glottis location and shape

Next, we investigated the properties of the latent space. We encoded all images of the BAGLS training dataset to gain a collection of latent space Ψ images. We first determined if any pixel is correlated for with the glottal area. We found that the correlation values follow a normal-like distribution centered around 0.00 with a standard deviation of 0.12 (Supplementary Figure **??**). We then investigated the value distribution of the latent space. The histogram shows a distribution centered around 0.8 (mean=0.75, median=0.78, mode=0.80), with a long tail towards 0 and a very short tail above 0.8 (Figure 2B). Interestingly, we clipped the available value space in the latent space between 0 and 6 (see Methods), however, the largest value we observed was 1.45, indicating that we are not constrained by our activation function.

**Fig 2.**
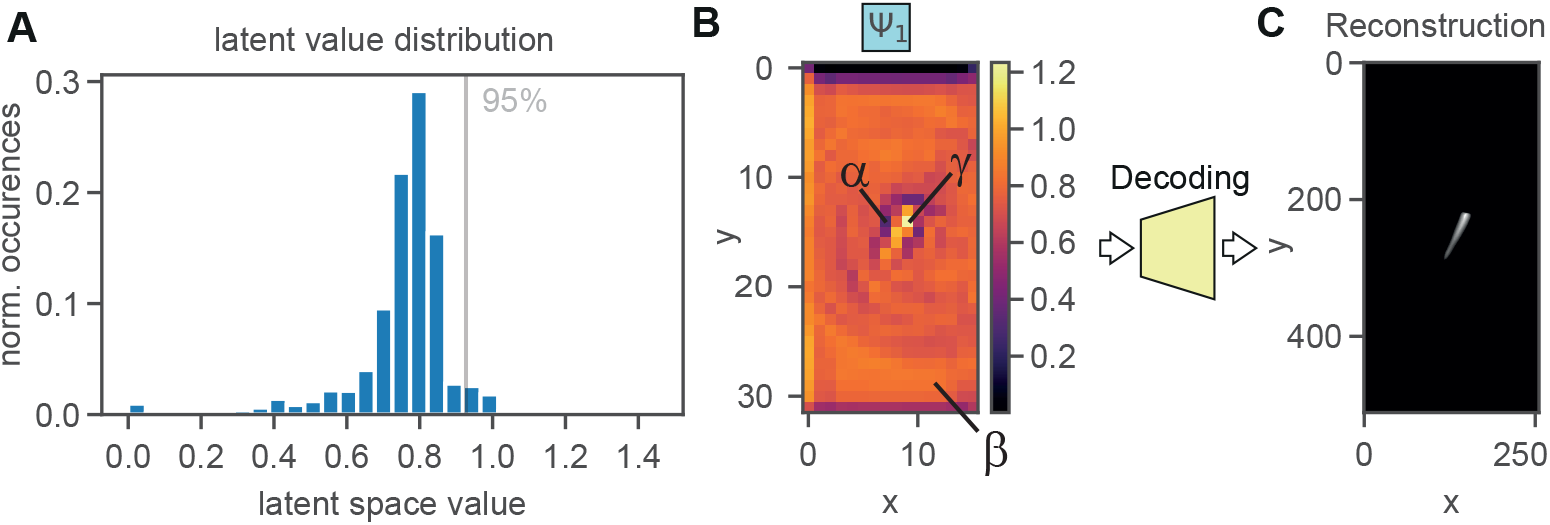
The latent space image Ψ_1_ values provide interpretable context. A: Value distribution of the latent space image Ψ_1_ across all pixels of all images in the BAGLS training dataset. We indicated the 95% confidence interval that is used for defining *γ* pixels. B: The average latent space image Ψ_1_ across 30 frames. We indicate the three pixel subtypes, *α* for glottis refining, *β* for background-defining, and *γ* for glottal area defining pixels. C: The average reconstruction obtained from feeding Ψ_1_ from panel B into the decoder.

To understand the meaning behind these values, we found that values around 0.8 seem to encode for background (referred to as *β* pixels), values higher than 0.8 define the glottal area (hereafter referred to as *γ*), and values lower of 0.8 are adjacent to glottal area-encoding values abbreviated *α* (Figure 2C). More examples are shown in Supplementary Figure **??**. Further, the latent space image Ψ_1_ is encoding the spatial location of the glottal area in *x* and *y*. We confirmed this by generating an artificial latent space image Ψ_1_ and varying *x* and *y* location, *γ* pixel value intensity and pixel drawing radius (Supplementary Movie 1). Values higher than 1.5 for *γ* resulted in image artefacts. We further investigated the role of the value drop (values *<* 0.8) for *α* pixels adjacent to the glottis-defining *β* pixels (values *>* 0.8). Our results suggest that *α* pixels in the surrounding of *γ* pixels are shaping the glottal area’s extent and refining its appearance (Supplementary Movie 2). Taken together, the interplay between *α* and *γ* pixels is crucial for an accurate glottis segmentation.

### Thresholded latent space is highly correlated with glottal area waveform

The glottal area waveform (GAW) is a time-variant signal important for assessing vocal fold physiology [2, 25]. We therefore asked, if the latent space image Ψ_1_ is a good proxy for the GAW. To answer this question, we used short video fragments from the BAGLS dataset and converted the provided ground-truth segmentation mask to the GAW (see Methods). We followed two approaches: (1) summing all values in Ψ_1_ and (2) threshold Ψ at 95% confidence interval (value = 0.8) and then summing the positive pixels. In Figure 3A, we show that approach (1) is correlated to a limited extend with GAWs (on average 0.03 ±0.56), however, approach (2) is highly correlated with the GAW, on average 0.84 ± 0.18. We show two exemplary videos with corresponding ground-truth GAW, the GAW generated by using the segmentation masks reconstructed by the decoder, and the thresholded Ψ_1_ waveform in Figure 3B.

**Fig 3.**
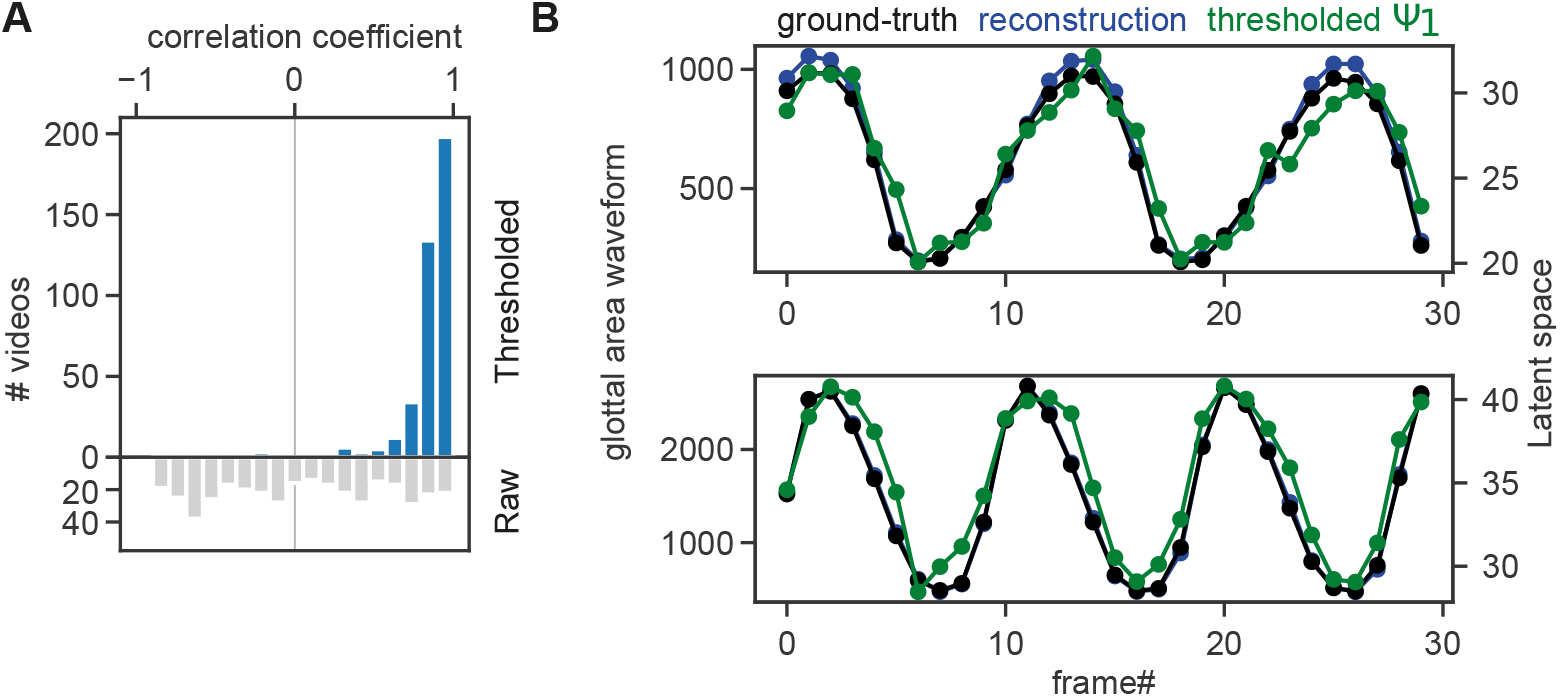
Thresholded latent space image Ψ_1_ is highly correlated with glottal area waveform. A: Distribution of correlation values across videos using either the raw latent space image Ψ_1_ or the thresholded Ψ_1_ using the 95% confidence interval as threshold (see Figure 2A). Correlation was computed from every latent space to glottal area waveform of 30 frames (N=399 videos). B: Two exemplary videos showing the original ground-truth glottal area waveform (GAW, black), the segmentation prediction of the deep neural network with a single latent channel (blue) and the thresholded latent space image Ψ_1_ from the same network (green).

### A low bit encoding is sufficient for glottis reconstruction

As the value range is very limited in the latent space (Figure 2B) and the existence of *α, β*, and *γ* pixels, we hypothesized that a low bit depth is sufficient for encoding the latent space Ψ_1_ for accurate glottal area reconstruction. By reducing the bit depth from 32-bit floating point to a range of 1 to 8-bit, we found that 4-bit encoding is sufficient for high quality reconstructions (Figure 4A-C). Specifically, with 4-bit encoding the IoU score becomes stable and shows a low error, that is neglectable with 8-bit encoding (Figure 4A). The mean-squared error (MSE) between full 32-bit reconstruction and low bit reconstruction declines as expected with increasing bit depth, but is still shows a significant deviation of from the original reconstruction (Figure 4B). We further are able to reproduce the high correlation of Ψ_1_ with the glottal area waveform (Figure 4D). In summary, we show that 4-bit encoding is sufficient for subjectively similar glottis segmentations compared to full 32-bit encoding.

**Fig 4.**
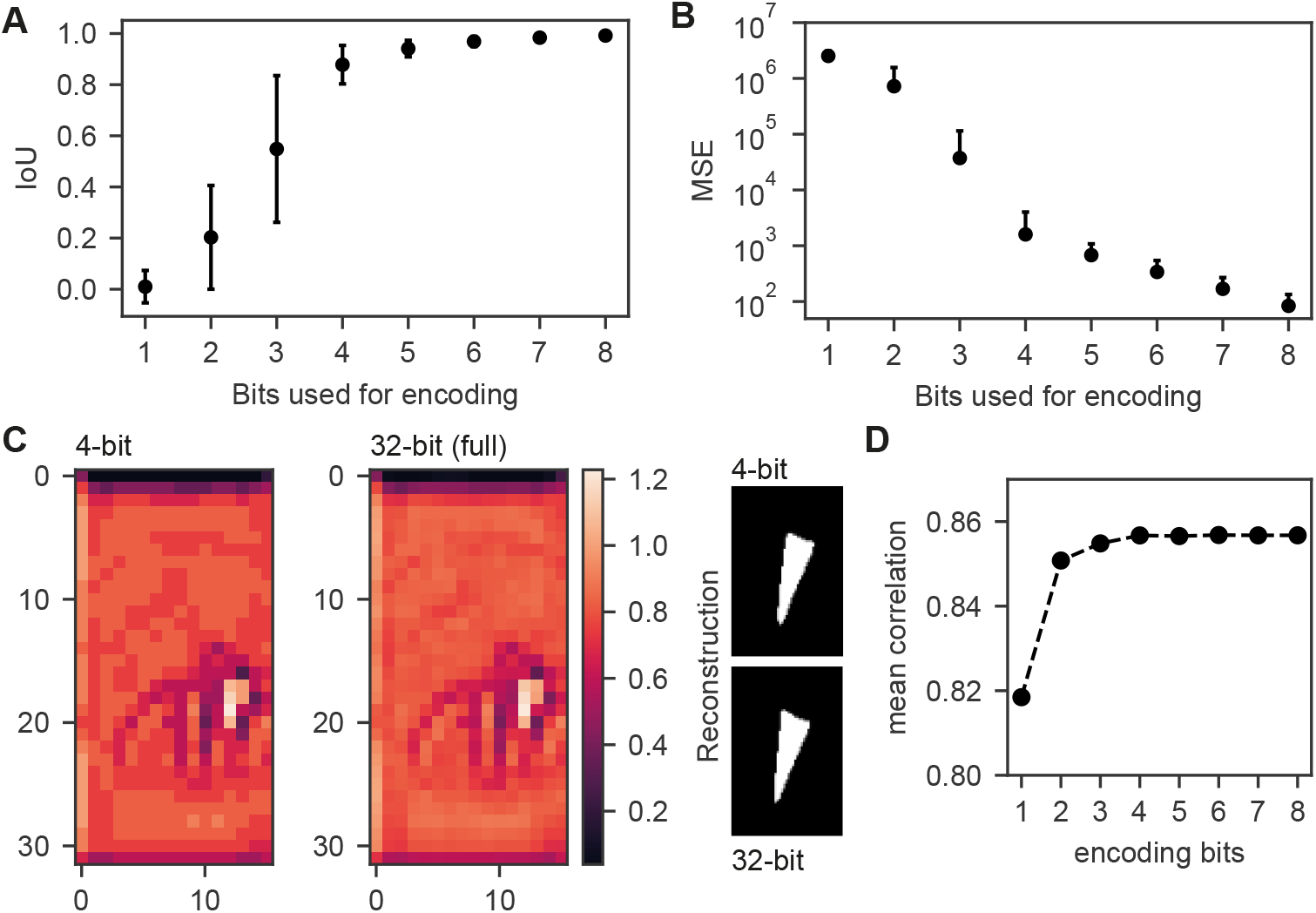
Four bits are sufficient for accurate reconstruction. A: IoU for reconstructions from lower bit encodings and full 32-bit reconstruction. B: Mean squared error (MSE) for lower bit reconstruction and full 32-bit reconstruction. C: Example for 4-bit and respective 32-bit encoding with respective reconstruction. D: Mean correlation of low bit latent space image Ψ_1_ and glottal area waveform (GAW).

### Light-weight decoders are capable of reconstructing the glottal area

As the latent space image Ψ is easily interpretable and shows a low-level complexity, we hypothesized that the decoder architecture can be largely simplified. Hence, we investigated how many convolutional filters and how many upsampling steps are necessary for decoding (Figure 5A). Further, we were interested if the upsampling strategy (nearest neighbours vs. bilinear interpolation) and multiple convolutional layers are affecting the decoding (Figure 5A). When using a single convolutional layer in each upsampling step, we found that one and two convolutional filters are not sufficient for decoding and that four convolutional filters are only sufficient in a single configuration (4x upsampling and bilinear interpolation) as shown in Figure 5B. The best results were achieved using eight convolutional filters together with 4x upsampling, which resulted in decent IoU scores (0.817, Figure 5B). Using two convolutional layers in each upsampling step, however, allowed 2x upsampling being competitive in the eight convolutional filters configuration. In general, two and four convolutional filters show better performance compared to the single convolutional layer experiment. However, these are not competitive with the configurations showing eight convolutional filters (Figure 5C). The top performance with two convolutional layers per block, eight convolutional filters and 4x upsampling with IoU=0.852 is slightly outperforming the single convolutional layer configurations. Despite the higher amount of trainable parameters in this configuration (Figure 5D), it has a relatively stable file size of 99 kB (Figure 5E). It is astonishing that even configurations with less than 200 trainable parameters achieve IoU scores higher than 0.4 (Table 1).

**Fig 5.**
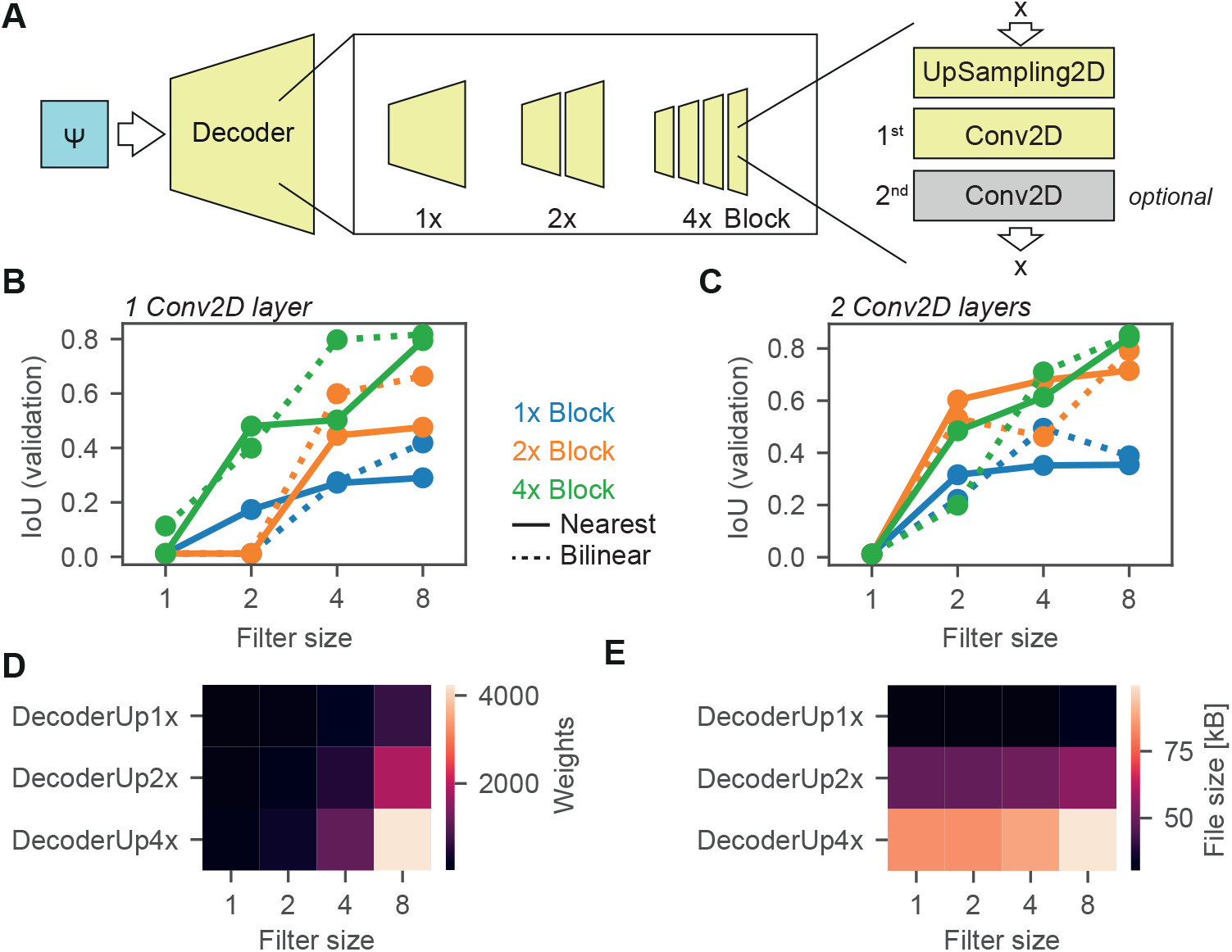
Lean decoders are sufficient for glottis reconstruction. A: Decoding from latent space. The decoder can consist of 1, 2 or 4 upsampling-convolution blocks, wherein one or two convolutional layers can be present. The filter size of each convolutional filter is fixed. B: IoU scores of different decoder blocks (color coded, 1 blue, 2 orange, 4 green) using either nearest neighbour (solid lines) or bilinear upsampling (dashed lines). C: Same as panel B, but with two convolutional layers. D: Trainable weights across decoder settings for two convolutonal layers per block. E: File size across decoder settings for two convolutional layers per block.

**Table 1.** Overview of all decoder configurations tested

| Blocks | Filter size | Conv2D layer | Weights | File size [kB] | IoU (nearest) | IoU (bilinear) |
| --- | --- | --- | --- | --- | --- | --- |
| 1x | 1 | One | 12 | 19 | 0.01 | 0.01 |
|  | 2 |  | 24 | 19 | 0.01 | 0.17 |
|  | 4 |  | 48 | 19 | 0.28 | 0.27 |
|  | 8 |  | 96 | 19 | 0.42 | 0.29 |
| 2x | 1 |  | 23 | 31 | 0.01 | 0.01 |
|  | 2 |  | 64 | 31 | 0.01 | 0.01 |
|  | 4 |  | 200 | 31 | 0.60 | 0.45 |
|  | 8 |  | 688 | 34 | 0.66 | 0.48 |
| 4x | 1 |  | 45 | 51 | 0.11 | 0.01 |
|  | 2 |  | 144 | 51 | 0.40 | 0.48 |
|  | 4 |  | 504 | 53 | 0.80 | 0.50 |
|  | 8 |  | 1872 | 58 | 0.82 | 0.79 |
| 1x | 1 | Two | 23 | 30 | 0.01 | 0.01 |
|  | 2 |  | 64 | 30 | 0.22 | 0.32 |
|  | 4 |  | 200 | 30 | 0.49 | 0.35 |
|  | 8 |  | 688 | 33 | 0.39 | 0.35 |
| 2x | 1 |  | 45 | 49 | 0.01 | 0.01 |
|  | 2 |  | 144 | 49 | 0.53 | 0.60 |
|  | 4 |  | 504 | 51 | 0.46 | 0.68 |
|  | 8 |  | 1872 | 55 | 0.79 | 0.72 |
| 4x | 1 |  | 89 | 84 | 0.01 | 0.01 |
|  | 2 |  | 304 | 84 | 0.20 | 0.48 |
|  | 4 |  | 1112 | 88 | 0.71 | 0.61 |
|  | 8 |  | 4240 | 100 | 0.85 | 0.84 |

## Discussion

In this work, we found that a single channel in the latent space of an encoder-decoder architecture is sufficient for glottal area reconstruction. We further show that the latent space forms an image that has interpretable properties, such as background (*β*), glottal area defining (*γ*) and refining (*α*) pixels. Our findings suggest that encoder-decoder frameworks are not only suitable for glottis segmentation, but also provide a higher-order approximation of the glottal area sufficiently encoded in a significantly smaller, single channel image. Together with a low bit encoding (Figure 4), it may serve as an efficient data storage system for glottis segmentations. The latent space image can be easily reconstructed using efficient decoders as presented in Figure 5.

In this study, we particularily focused on the latent space and its sufficiency for glottis segmentation. We found that removing the U-Net-specific skip connections yielded lower IoU scores in the validation set, whereas we did not find any differences in the test set, i.e. on independent, unseen data (Figure 1D). This is in line with a previous study [10], where the authors showed that the kind of skip connection is not important (adding or concatenating channels from encoder to decoder). They also found a significant drop in the validation IoU score, when removing the skip connections. However, we have not specifically investigated the role of skip connections in this context. It remains elusive if certain data configurations benefit from the skip connections.

Glottis segmentation is a straight forward task and was previously approached using thresholding-based techniques [7, 20, 26]. Therefore, it is likely that the encoder-decoder architecture learns a non-linear thresholding algorithm. However, other modalities, such as anterior-posterior point prediction for midline estimation [19] and vocal fold localization for paralysis analysis [27] may not benefit from this very constrained latent space. Future studies should address these limitations and speculate about the necessity of an increased latent space crucial for multitask architectures, as the latent space has been shown useful for midline estimation [19].

The U-Net is a very powerful starting point for biomedical image segmentation tasks, also for glottis segmentation [4, 9, 28]. Modifications to this architecture, such as convolutional layers with LSTM memory cells [29] as shown by [28] may improve the glottal segmentation accuracy. Also, more sophisticated encoding backbones, such as the ResNet [30] and the EfficientNet [31] architecture show superior performance in glottis segmentation, especially on more dissimilar data sources [20]. Future research should investigate, if these architectures are able to detect and encode better high-level features in the latent space, such that a potentially higher dimensionality in the latent spcae yields further performance improvements.

## Conclusion

With this work, we contribute to the understanding how glottis segmentation is performed by deep neural networks and that we are able to uncover the deep neural network black box character by identifying three value ranges with a specific role, namely *α, β* and *γ* pixels. Future studies may elucidate if these three subclasses can be further refined and if they occur across architectures and segmentation tasks. In general, our findings also allow very efficient architectures leveraging the potential of real-time applications of glottis segmentations in a clinical setting and maybe used together with recent advances in open HSV systems [32]. Further research on quantitative measures may include how the latent space image Ψ influences these computations and if the latent space is also sufficient for approximating complex quantitative parameters to assess easily voice physiology.

## Supporting information

**S1 Fig. Latent space pixels are not directly correlated with glottal area**.

**S2 Fig. Latent space coding is consistent across recordings**.

**S1 Video**. *γ* **pixels define glottal area** The glottal area is defined in a range between 0.8 and 1 leading to a nice segmentation. Values higher than 1 tend to create artefacts.

**S2 Video**. *α* **pixels shape glottal area**. This movie shows the interplay between *α* and *γ* pixels.

## Acknowledgments

This project was supported by Deutsche Forschungsgemeinschaft (DFG) under grant no. SCHU 3441/3-2.

## Author contributions

AMK conceived the project, trained deep neural networks, analyzed the data, created figures and tables. AS, SD and MS interpreted data. AS secured funding. AMK wrote the paper with input from AS, MS and DS.

## Notes

https://github.com/ankilab/latent

https://zenodo.org/record/5772799

### Competing Interest Statement

The authors have declared no competing interest.

